# Long-Term Persistence of Plasmids Targeted by CRISPR Interference in Bacterial Populations

**DOI:** 10.1101/2021.04.03.438301

**Authors:** Viktor Mamontov, Alexander Martynov, Natalia Morozova, Anton Bukatin, Dmitry B. Staroverov, Konstantin A. Lukyanov, Yaroslav Ispolatov, Ekaterina Semenova, Konstantin Severinov

## Abstract

CRISPR-Cas systems provide prokaryotes with an RNA-guided defense against foreign mobile genetic elements (MGEs) such as plasmids and viruses. A common mechanism by which MGEs avoid interference by CRISPR consists of acquisition of escape mutations in regions targeted by CRISPR. Here, using microbiological, live microscopy, and microfluidics analyses we demonstrated that plasmids can persist in *Escherichia coli* cells at conditions of continuous targeting by the type I-E CRISPR-Cas system without acquiring any genetic alterations. We used mathematical modeling to show how plasmid persistence in a subpopulation of cells mounting CRISPR interference is achieved due to the stochastic nature of CRISPR interference and plasmid replication events. We hypothesize that the observed complex dynamics provides bacterial populations with long-term benefits due to the presence of mobile genetic elements in some cells, leading to diversification of phenotypes in the entire community and allowing rapid changes in the population structure to meet the demands of a changing environment.

## Introduction

CRISPR (Clustered Regularly Interspaced Short Palindromic Repeats)-Cas (CRISPR associated genes) is a widespread form of adaptive immunity in prokaryotes^1^. CRISPR-Cas systems are able to recognize and destroy nucleic acids with sequences complementary to spacers stored in CRISPR arrays^2,3,4^. In the array, spacers are separated by the repeat sequences. CRISPR array transcripts are processed into individual CRISPR RNAs (crRNAs) containing spacer sequences with flanking repeat fragments. Individual crRNAs bind to Cas proteins forming an effector complex, which can recognize protospacers — sequences complementary to the crRNA spacer part. For CRISPR-Cas effectors targeting DNA, the recognition requires, in addition to full or partial complementarity between crRNA spacer and the protospacer, the presence of PAM, a protospacer adjacent motif, that is recognized by the protein part of the effector complex^5^

Multiple examples of protection of prokaryotic cells by different CRISPR-Cas systems acting through the CRISPR interference mechanism described above from infection by DNA and RNA viruses and transformation by plasmids have been documented^6,7,8^. Depending on the virus and the type of CRISPR-Cas system, a cell mounting the interference response can clear the infection and survive or die in the course of abortive infection. In the latter case, the population as a whole benefits because the appearance of progeny viruses is prevented^9^. Viruses respond to the pressure from CRISPR-Cas by acquiring point mutations in protospacers targeted by crRNAs or in their PAMs^10^. In turn, cells respond to such viral escapers by updating their CRISPR memory by acquiring additional viral-derived spacers^11^.

During plasmid transformation/conjugation experiments CRISPR interference results in decreased efficiencies of DNA uptake. Mutations in targeted protospacers or their PAMs restore transformation efficiencies^6^. In experiments where cells are forced to keep a plasmid targeted by CRISPR-Cas by inclusion of an appropriate antibiotic in the medium, mutations inactivating CRISPR-Cas system components are observed^12,13,14^.

Given a considerable interest in potential use of CRISPR-Cas targeting of antibiotic-resistance plasmids as means to reduce antibiotic resistance spread, we here undertook a study of the interaction of the well-studied *Escherichia coli* type I-E CRISPR-Cas system^11,15,16^ with plasmids carrying protospacers recognized by the Cascade effector complex. We were specifically interested in colonies formed on antibiotic-containing selective media by cells with an active CRISPR-Cas system transformed with plasmids carrying protospacers targeted by the effector. We report that only a small fraction of resulting colonies are formed by cells with inactivated CRISPR-Cas. Most colonies have an active CRISPR-Cas system and unaltered plasmids which are subject to CRISPR interference. Using a combination of microbiological, microscopic and microfluidics experiments we show that cells in such colonies are heterogeneous, with most cells having little or no plasmid. Apparently, these colonies are formed due to the presence of a minor fraction of cells that manage to keep the plasmid at conditions of ongoing CRISPR interference. We use mathematical modeling to show how plasmids stably persist in generations of such cells due to a balance of CRISPR interference and plasmid replication rates.

Our results show that potentially beneficial plasmids can be stably maintained in bacterial populations while being targeted by CRISPR, allowing rapid expansion of plasmid-bearing subpopulations when conditions demand.

## Results

### Colonies formed after transformation of protospacer plasmids into cells mounting CRISPR interference contain cells with active CRISPR-Cas and unchanged plasmids

The *Escherichia coli* KD263 cells that contain inducible *cas* genes and a CRISPR array with a single g8 spacer^17,18^ were grown in the presence or in the absence of *cas* gene expression inducers and transformed with pG8, a pUC-based plasmid containing the g8 protospacer with an interference-proficient ATG PAM^11^. After transformation, cells without *cas* gene induction (CRISPR OFF) were plated on a medium supplemented with ampicillin to select colonies formed by plasmid-bearing cells. Pre-induced CRISPR ON cells were plated on a medium that contained both ampicillin and inducers of *cas* genes expression (Fig. 1a). Compared to CRISPR OFF cells, approximately 200 times less ampicillin-resistant colonies were formed after the transformation of CRISPR ON cells (Fig. 1b). No difference in the number of transformants was observed when a plasmid vector without the g8 protospacer was used for transformation, indicating that the drop in transformation efficiency was due to CRISPR interference mounted when the g8 protospacer in pG8 was recognized by the Cascade effector charged with the g8 crRNA. A similar experiment with another plasmid, pRSFG8, which provides resistance to kanamycin, showed similar results: ~50-80 times less transformant colonies were formed by CRISPR ON cells compared to CRISPR OFF cells (Fig. 1b). With both plasmids, antibiotic-resistant colonies formed by CRISPR ON cells were visually indistinguishable from CRISPR OFF cells colonies.

**Figure 1.**
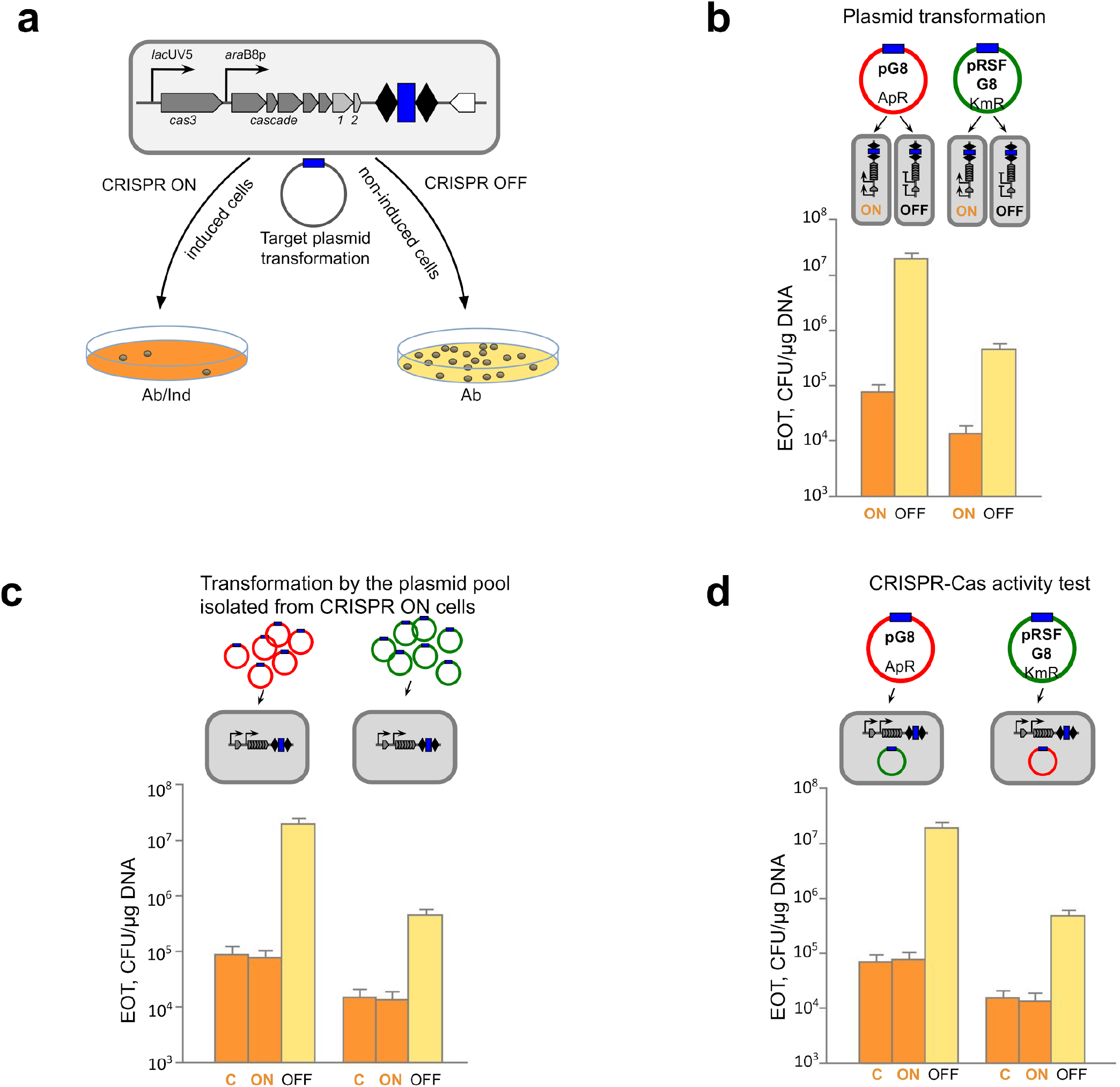
Cells forming colonies on selective media under CRISPR ON conditions contain plasmids that are subject to interference and a functional CRISPR-Cas system. **a,** An *E. coli* KD263 cell harboring *cas* genes controlled by inducible promoters and a CRISPR array with two repeats (black rhombi) and a single g8 spacer (blue rectangle) is schematically shown at the top. Cells are grown in the presence or in the absence of *cas* gene expression inducers to prepare, correspondingly, CRISPR ON and CRISPR OFF competent cells, which are transformed with a plasmid bearing the g8 protospacer (shown as a blue rectangle on a circle representing the plasmid, fully matches the g8 spacer) with a functional PAM. After transformation, CRISPR ON cells are plated on a medium supplemented with *cas* genes inducers and an appropriate antibiotic (Ab/Ind); CRISPR OFF cells are plated on a medium containing only the antibiotic (Ab). **b,** CRISPR ON and CRISPR OFF cells were transformed with ampicillin-resistant pG8 or kanamycin-resistant pRSFG8 plasmids bearing the g8 protospacer (blue rectangle) and efficiency of transformation (EOT) was determined as CFUs per μg of plasmid DNA. Bars show mean EOTs from three independent experiments. Standard deviations of the mean are indicated. **c,** Plasmids purified from CRISPR ON colonies transformed with either pG8 or pRSFG8 were retransformed into CRISPR ON or CRISPR OFF competent cells and EOT was determined. Transformation of CRISPR ON cells with initial pG8 and pRSFG8 plasmids was used as a control (“C”). Bars show mean EOTs from three independent experiments. Standard deviations of the mean are indicated. **d,** Competent cells prepared from cells from CRISPR ON colonies transformed with pG8 or pRSFG8 were transformed with compatible g8 protospacer plasmids (cells bearing pG8 were transformed with pRSFG8 and vice versa). As a control, transformation of plasmid-less CRISPR ON cells with pG8 and pRSFG8 plasmids was performed (“C”). Bars show mean EOTs from three independent experiments. Standard deviations of the mean are indicated.

To test whether plasmids in CRISPR ON colonies escaped interference by accumulating mutations, plasmids from ten randomly chosen individual CRISPR ON colonies were purified and retransformed into CRISPR ON and CRISPR OFF competent cells. In every case, a drop in transformation efficiency into induced cells was the same as that observed during the original transformation experiment (Fig. 1c). We therefore conclude that plasmids present in CRISPR ON colonies are subject to interference by CRISPR effector charged with g8 spacer crRNA and in this respect are identical to plasmids used in the original experiment. Consistently, sequencing of the protospacer region from plasmid prepared from pooled CRISPR ON colonies did not reveal differences from the pG8 sequence (Extended Data Fig. 1).

To determine whether CRISPR ON cells forming colonies on selective medium contain a functional CRISPR-Cas system, competent cells were prepared from CRISPR ON transformants and transformed with compatible plasmids carrying the g8 protospacer and a different antibiotic resistance marker. Cells derived from pG8-transformed CRISPR ON colonies interfered with pRSFG8 transformation as efficiently as induced control plasmidless KD263 cells (Fig. 1d). The same situation was observed when competent cells derived from pRSFG8-transformed CRISPR ON colonies were transformed with pG8 (Fig. 1d). We therefore conclude that CRISPR ON colonies transformed with plasmids carrying a protospacer matching crRNA spacer are formed by cells with a functional CRISPR-Cas system.

### Cells from CRISPR ON colonies contain less plasmids than CRISPR OFF colony cells

Quantitative PCR with plasmid-specific primers was used to determine the plasmid copy number (PCN) in cells from CRISPR ON and CRISPR OFF colonies. Amplification of the chromosomal *gyrA* gene was used for normalization (see Methods). On average, there were 233 ± 46 copies of pG8 per cell in CRISPR OFF colonies (Fig. 2b, top row panel), which is consistent with PCN values for the pUC vector on which pG8 is based^19^. For pRSFG8, an average value of 119 ± 21 copies per CRISPR OFF colony cell was calculated (Fig. 2c, top row panel), which is also consistent with published data^19,20^. In contrast, cells from CRISPR ON colonies had an average PCN of 0.18 ± 0.06 (for pG8) and 0.71 ± 0.27 (for pRSFG8) (Figs. 2b and 2c, top row panels).

**Figure 2.**
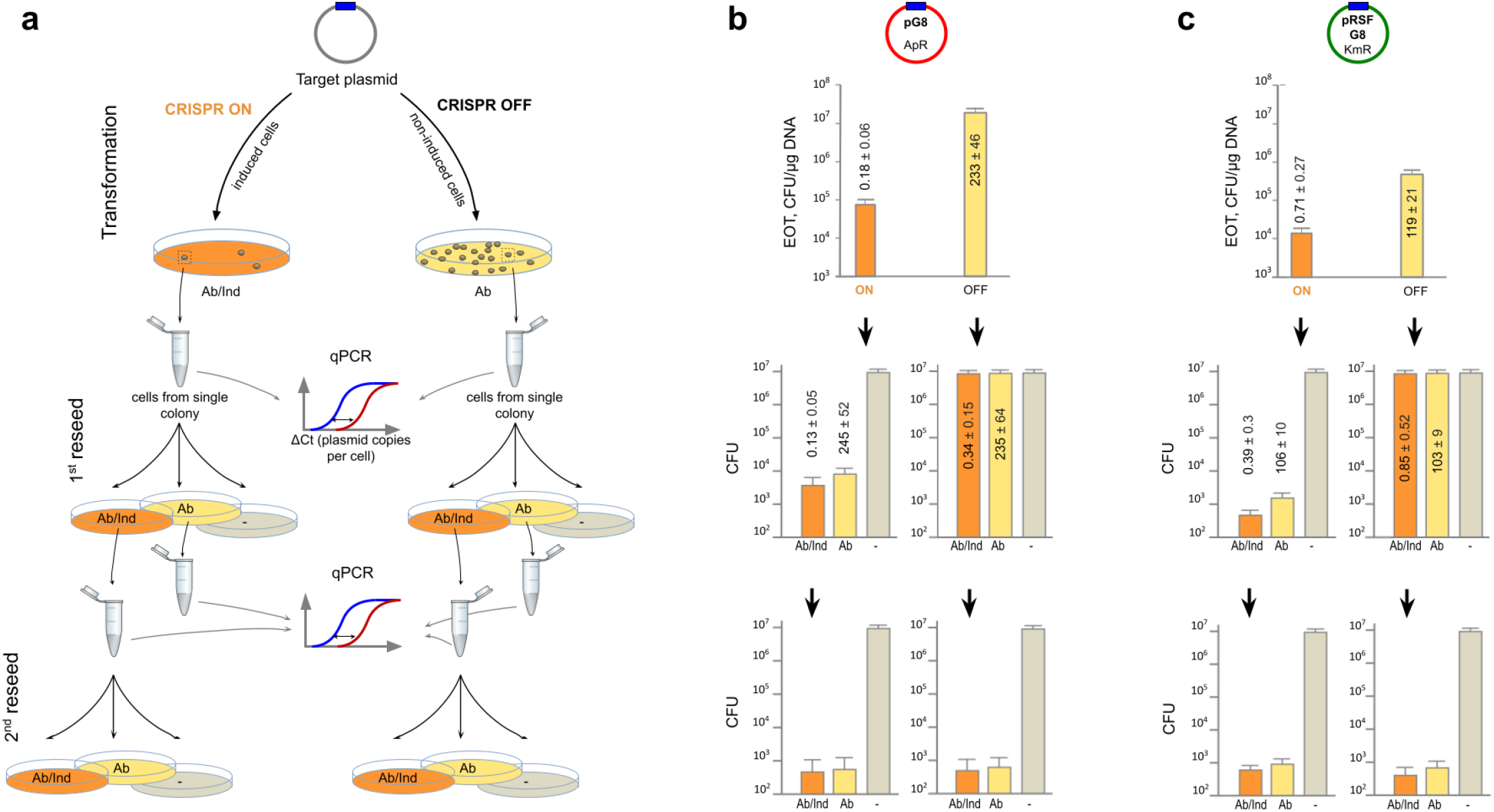
Colonies formed at CRISPR ON conditions mostly contain plasmid-less cells. **a,** Cells from CRISPR ON and CRISPR OFF colonies obtained as in **Fig. 1** are reseeded on media supplemented with antibiotic and inducers (“Ab/Ind”, orange), antibiotic only (“Ab”, yellow), or plates with neither inducers nor antibiotics (“-”, grey). Cells from colonies formed on Ab/Ind plates are reseeded the second time. Real-time PCR is used to determine PCN per cell in colonies formed at each condition. **b** and **c,** Experiment was done as outlined in panel **a** using the pG8 and pRSFG8 plasmids, correspondingly. Numbers show mean PCN values obtained from three independent qPCR experiments with different colonies. Standard deviations of the mean are indicated.

The below 1 PCN value indicates that many cells in CRISPR ON colonies are plasmid-less and the colonies should thus be heterogeneous. To determine the ratio of plasmid-bearing and plasmid-less subpopulations we replated cells from randomly chosen CRISPR ON and CRISPR OFF colonies transformed with pG8 or pRSFG8 on three types of media (Fig. 2a). Plating on a medium with no *cas* gene expression inducers and without an antibiotic determined the total number of viable cells. Plating on a medium supplemented with an appropriate antibiotic determined the number of viable plasmid-bearing cells. Plating on a medium supplemented with *cas* gene expression inducers and an appropriate antibiotic allowed us to determine whether cells from CRISPR ON colonies that carried a plasmid were losing it during growth under conditions of continued CRISPR interference.

For CRISPR OFF transformants, the number of colonies formed upon reseeding on plates with and without antibiotics was the same (Figs. 2b and 2c, right panels in the middle row) indicating that both pG8 and pRSFG8 are stably maintained in the absence of antibiotic selection. In contrast, only one out of a few thousands reseeded cells from CRISPR ON colonies grew on antibiotic-containing plates (Figs. 2b and 2c, left panels in the middle row). Thus, most cells in CRISPR ON colonies indeed lost the plasmid and must have survived due to the presence of a minor fraction of plasmid-bearing cells that decreased antibiotic concentration within the colony.

The number of colonies formed by cells from CRISPR ON colonies on plates supplemented with both *cas* gene expression inducers and an antibiotic was further decreased 5-10-fold compared to the number of colonies grown on plates with antibiotic only (compare Figs. 2b and 2c, left panels in the middle row). This indicates that CRISPR interference continues to purge plasmids from CRISPR ON plasmid-bearing cells, albeit at an efficiency that is considerably lower than that observed during plasmid transformation.

The number of colonies observed after reseeding cells from CRISPR OFF colonies on a medium containing both the *cas* genes inducers and an antibiotic was the same as that on the medium with antibiotic only or without any additions (Figs. 2b and 2c, right panels in the middle row). This result seems to indicate that interference against an established plasmid is inefficient. Yet, quantitative PCR showed that per cell PCN values for colonies formed upon reseeding of original CRISPR OFF colonies on media with *cas* genes inducers and an antibiotic were as low as those for initial CRISPR ON transformants (Figs. 2b and 2c, left panels in the middle row). In contrast, PCN per cell for colonies formed on plates containing antibiotic only was as high as in corresponding CRISPR OFF colonies, implying that PCN restores to normal levels in the absence of CRISPR interference.

The second round of reseeding confirmed that most cells in colonies originating from CRISPR OFF colonies have lost plasmids after growth at conditions of induction of *cas* genes expression (Figs. 2b and 2c, right panels in the bottom row). In addition, the proportion of plasmid-bearing cells was further decreased when cells from a CRISPR ON colony formed after the first reseed were replated in the presence of *cas* genes inducers and an antibiotic. The effect was ~100 fold for pG8 colonies and less pronounced for pRSFG8 colonies (compare Figs. 2b and 2c, left panels in the bottom row).

### Direct observation of plasmid-bearing cells in CRISPR ON colonies

Plasmid pG8-GFP, a derivative of pG8 carrying a constitutively expressed green fluorescent protein TagGFP2 gene, was created to allow direct observation of plasmid-bearing cells. Similarly to pG8, the pG8-GFP plasmid was subject to CRISPR interference as evidenced by a ~200-fold decrease in the number of colonies formed at CRISPR ON conditions compared to CRISPR OFF conditions. While all CRISPR OFF colonies transformed with pG8-GFP were highly fluorescent when irradiated with a handheld UV lamp, most CRISPR ON colonies were dim. Only 1-3% of all CRISPR ON colonies fluoresced (Fig. 3a). Retransformation experiments conducted as in Fig. 1c revealed that plasmids from these rare colonies did not contain escape mutations. However, whole genome sequencing of DNA extracted from two randomly chosen fluorescent CRISPR ON colonies revealed frame-shift mutations in the *cse1* gene and/or in the *araBp8* promoter from which the *cas* operon is transcribed (Extended Data Table 1).

**Figure 3.**
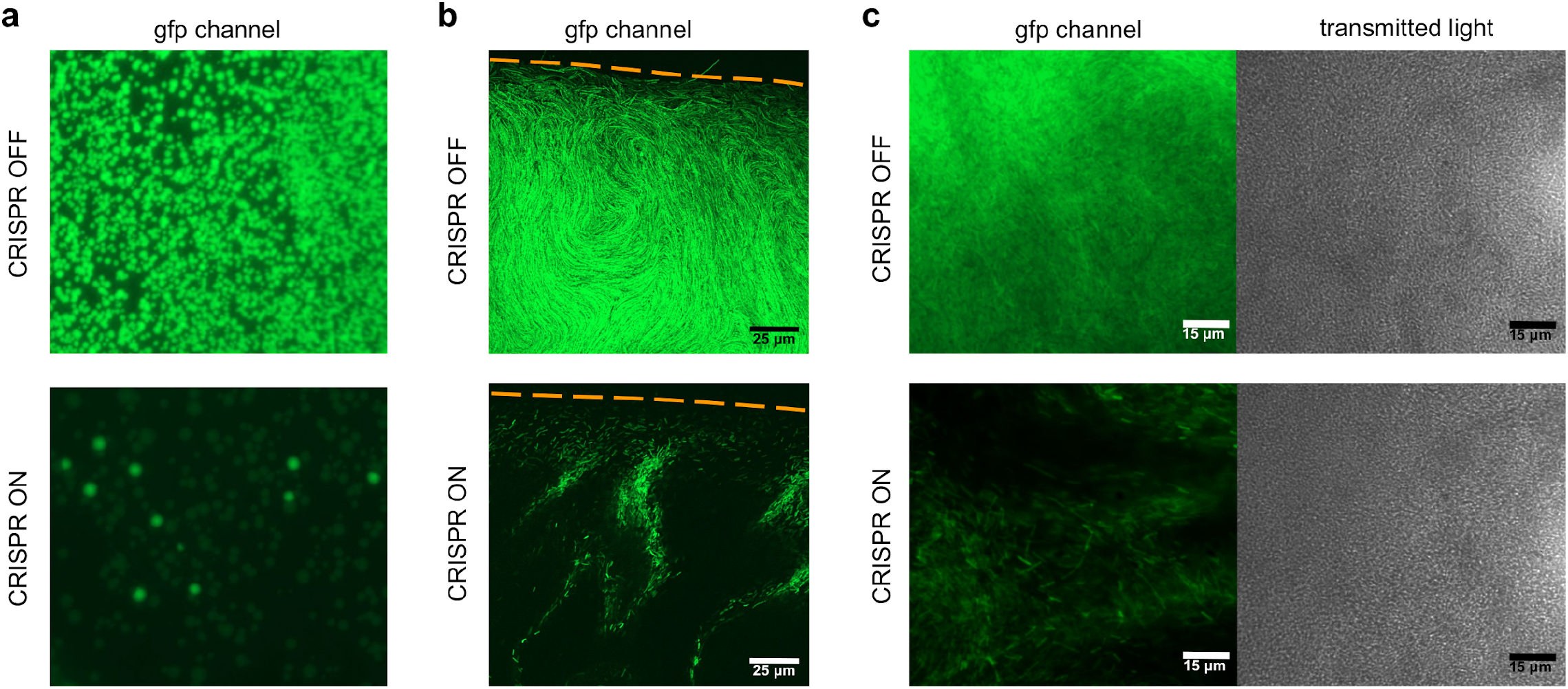
Fluorescence microscopy of *E. coli* KD263 colonies. **a,** Images of fragments of plates containing CRISPR ON and CRISPR OFF transformant colonies. Note that the majority of CRISPR ON colonies are weakly fluorescent; their internal structure at a higher magnification is shown in panels **b** and **c**. Highly fluorescent CRISPR ON colonies have a non-functional CRISPR-Cas system (see Extended Data Table 1). **b,** Images of a CRISPR OFF and a dim CRISPR ON colony periphery obtained using confocal microscopy in the gfp channel. Orange dashed lines show colony edges. **c,** Wide-field fluorescence microscopy of CRISPR OFF and dim CRISPR ON colonies.

The individual dim CRISPR ON and CRISPR OFF colonies were analysed using confocal and wide-field fluorescence microscopy at high magnification (Fig. 3b,c, Extended Data Fig. 2). In the CRISPR OFF colonies, all cells were brightly fluorescent. In contrast, only a small fraction of cells possessed detectable fluorescence in the CRISPR ON colonies. Radially extended serpentine-shaped rows of fluorescent cells on the background of non-fluorescent plasmid-less cells that are clearly seen at the colony edges are consistent with inherited maintenance of plasmids in some lineages within the colony. The absence of expanding fluorescent sectors seems to suggest that most cells in such lineages lose the plasmid with time (Fig. 3b).

Our results show that most CRISPR ON colonies (each derived from a single founder cell transformed with the plasmid bearing the target protospacer) are heterogeneous and most cells in such colonies are either completely or nearly plasmid-less. Clearly, for continued colony growth on selective medium there must be at least one uninterrupted line of plasmid-bearing cells that persists through multiple generations. To study the distribution of plasmid-bearing fluorescent cells we analysed cells from dim CRISPR ON colonies transformed with pG8-GFP using flow cytometry (Fig. 4a). Cells transformed with the pG8 plasmid bearing no fluorescent marker were used to define the level of autofluorescence. Cells derived from pG8-GFP transformed CRISPR OFF colonies were used as a positive control. The results showed that about 10% of cells from CRISPR ON colonies were fluorescent (Figs. 4b,c). However, the mean intensity of fluorescence of such cells was ~7 times less than that of positive control cells. Detailed statistics demonstrated that PCN (assumed here to be directly proportional to fluorescence) in plasmid-bearing fraction of CRISPR ON cells reached an upper limit corresponding to 0.3-0.4 of that in CRISPR OFF cells (Fig. 4c).

**Figure 4.**
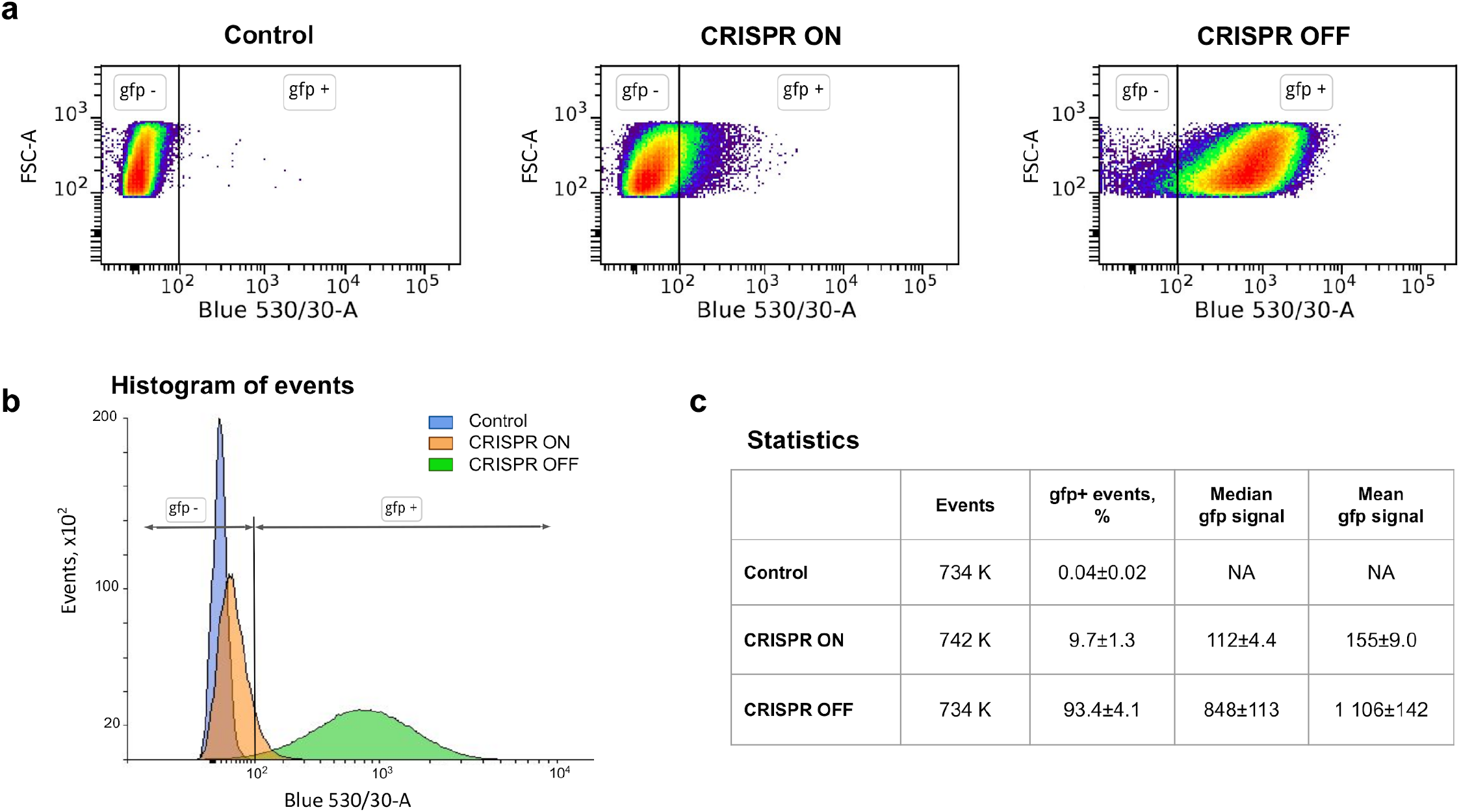
Flow cytometry analysis of cells from CRISPR ON and CRISPR OFF colonies. **a,** Density plots in forward scattering (FSC-A) and green fluorescence (Blue 530/30-A) channels. Vertical black lines in all panels show borders between gfp-gate consisting of non-fluorescent events and gfp+ gate consisting of fluorescent events. The control panel shows cytometry results of cells bearing the pG8 plasmid (without GFP) as a negative control. **b,** Histogram of the distribution of fluorescence levels in cells from indicated colonies. **c,** Statistics of flow cytometry results.

### Direct real time observation of plasmid loss due to CRISPR interference

To observe plasmid loss caused by CRISPR interference in real time, cells from CRISPR OFF colonies transformed with pG8-GFP were used to seed microfluidic growth chambers (Extended Data Fig. 3). Growth chambers seeded with single cells were observed for 7 hours in the presence or in the absence of *cas* gene expression inducers in the medium flowing through the main channel of the microfluidic device. No antibiotic was added. As expected, in the absence of *cas* genes expression inducers, the founder cells divided and all progeny remained highly fluorescent (Fig. 5a, Extended Data Figs. 4,5). In contrast, in the presence of inducers, progeny cells remained fluorescent only until the fifth division (Fig. 5b). Afterwards, most cells ceased to fluoresce, presumably due to the earlier loss of plasmid caused by CRISPR interference and dilution of the GFP protein present in the founder cell and its immediate descendants. However, some cells retained fluorescence. Though some descendants of such fluorescent cells subsequently lost fluorescence, others formed lineages of fluorescent cells that persisted during the time of the experiment. The schematic tree of cell divisions illustrates that one single cell is able to generate a branch of fluorescent cells, as well as multiple branches of offspring that lost fluorescence (Fig. 5c).

**Figure 5.**
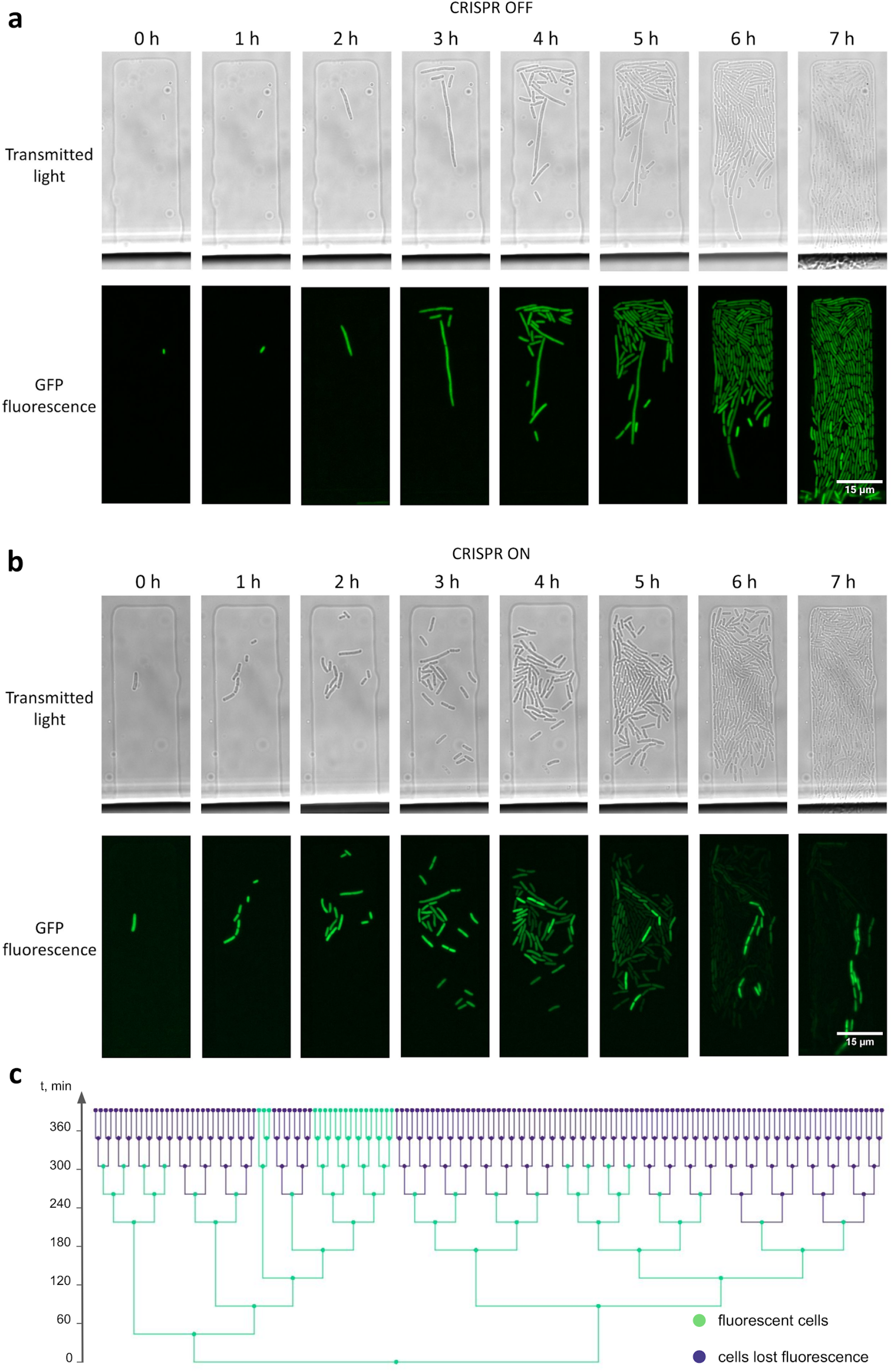
Live fluorescence microscopy of cells in a microfluidics device. **a,** The divisions of a single CRISPR OFF cell bearing the pG8-GFP plasmid over time in a microfluidics chamber supplemented with LB medium. **b,** As in **a** but in a medium containing *cas* gene expression inducers. **c,** A tree of cell divisions depicts the loss of fluorescence through cell generations in a microfluidics chamber shown in panel **b**. The length of the branches schematically illustrates the time between subsequent divisions.

## Discussion and theoretical analysis

In this work, we show that plasmids can survive in cells mounting an active CRISPR interference response against them. We experimentally ruled out trivial possibilities such as inactivation of CRISPR-Cas system or accumulation of escape mutations in such cells. While only a small fraction of cells retain plasmids under the ongoing pressure from the CRISPR-Cas system, plasmids remain in these cells and their descendants for many generations, such that apparently healthy colonies are formed on selective media containing antibiotics in concentrations sufficient to completely prevent growth of cells without plasmids.

Plasmids providing resistance to ampicillin rely on β-lactamase secreted in the periplasm to degrade the antibiotic outside the cell^21,22^. The phenomenon of indirect resistance where ampicillin-resistant colonies decrease the concentration of ampicillin in the medium and support the growth of susceptible satellite colonies is well known^23,24^. In our experiments with pG8 and pG8-GFP plasmids, such indirect resistance is apparently responsible for the observed CRISPR ON colonies heterogeneity with a small number of plasmid-bearing cells supporting the growth of a much larger number of cells that have lost the plasmid. At the same time, previous studies demonstrated the absence of indirect resistance for kanamycin^24^. Thus, the mechanism of growth of CRISPR ON colonies with pRSFG8 remains unclear and may involve specific three-dimensional arrangement of cells within a colony^23,25^.

While all our experiments clearly demonstrate that the majority of cells in CRISPR ON colonies consist of cells completely devoid of plasmids and the average PCN in rare plasmid-bearing cells is fewfold less that that in CRiSPR OFF cells, the measurements made by replating, flow cytometry, and in microfluidic device agree with each other only quantitatively. The possible reasons for such discrepancies could lie in difference in time elapsed after transformation, varying levels of *cas* gene induction, and other more specific distinctions between experimental setups. These differences notwithstanding, we below suggest an explanation to the apparently probabilistic and history-dependent response of PCN to CRISPR interference based on a simple stochastic model of plasmid replication and interference. For multicopy plasmids used here, when only one or a few plasmids are present in the cell, both the interference and replication kinetics should be limited by plasmid concentration, so the per plasmid rates of both processes are constant, i.e. independent of PCN. This means that per population rates of both processes depend linearly on the number of plasmids present in the cell. Yet when the PCN is large and close to the stationary number of plasmids [Pl]_st_, the replication rate should approach zero. Similarly, for all reasonable forms of CRISPR interference kinetics, an increase in the number of plasmids should result in a progressively smaller increase in the interference rate and its eventual saturation to a constant when the concentration of plasmids becomes high. Three scenarios that satisfy these general constraints are possible:

- The replication rate is always lower than the interference rate (Fig. 6a).
- The replication rate is higher than the interference rate until a certain threshold plasmid copy number is reached (Fig. 6b).
- There is a range of PCN values for which the replication rate exceeds the interference rate; beyond this range the interference rate is higher (Fig. 6c).

**Figure 6.**
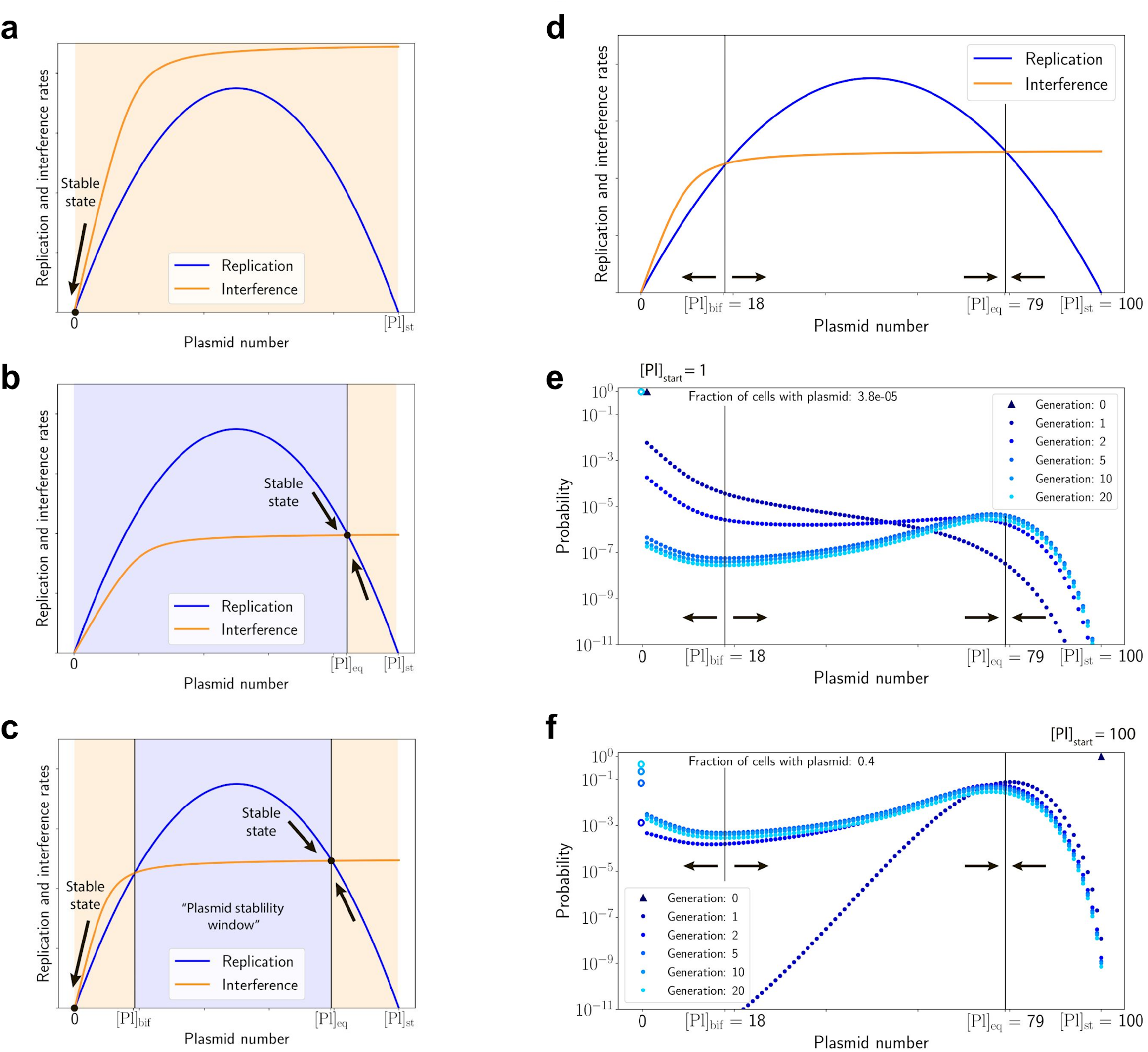
Plasmid copy number dynamics in a CRISPR ON cell. **a-c,** Three possible scenarios between plasmid replication (blue lines) and CRISPR interference (orange lines) rates: **a,** The interference rate is higher than the replication rate for any PCN, plasmids quickly become extinct. **b,** The replication rate is higher than the interference rate, PCN quickly reaches an equilibrium point [*Pl*]_eq_. **c,** There exists an intermediate range of PCN values, [*Pl*]_bif_ < [*Pl*] < [*Pl*]_eq_, where the replication rate is higher than the interference rate; beyond this range the interference dominates. Ranges of PCN where replication or interference rates dominate are shown by blue or orange shading, respectively. **d,** The replication (blue line) and interference (orange line) rates used in the solution of the master equation (5), parametrized as the Logistic (Eq. (1)) and Michaelis-Menten (Eq. (3)) kinetics. **e,** The probability P_n_ (t) for a cell to have n plasmids just before their partition between two daughter cells after 1,2, 5, 10, and 20 generations shown by dots of varying shades of blue. Initially, a single plasmid was introduced into a cell, which is marked by a triangle in the upper left corner. Empty circles, also marked by shades of blue of the corresponding generations, show the fraction of cells that lost all plasmids. **f,** Same as in **e**, but for the initial number of plasmids equal to [Pl]_st_, marked by a triangle in the upper right corner. After ≈ 10 generations, the probability distribution P_n_(t) converges to the universal form, shown by a pale blue line in panels (**e**) and (**f**).The parameters used in these solutions are listed in the Methods section. Vertical black lines in panels **d-f** show stable and unstable fixed points of PCN dynamics, their stability is shown by converging and diverging arrows.

The first scenario leads to quick loss of plasmids in all cells; the second results in survival of plasmids in the majority of cells at an equilibrium PCN [Pl]_eq_ value that is less than that in the absence of interference. The third scenario explains prolonged survival of plasmids in a small fraction of cells observed in our experiments. Since all transformed cells initially have just one copy of a plasmid, most lose it since at low PCN the interference rate is higher than the replication rate. However, due to an intrinsic stochasticity of interference and replication events, there is a small but finite possibility that in some transformed cells plasmid replication events occur more often than interference events. If such a favourable (for plasmid) situation occurs, the PCN may go over a “bifurcation threshold” (marked as [Pl]_bif_ in Fig. 6c), above which the replication rate exceeds the interference rate. From this point on plasmids will likely survive and continue to expand deterministically until reaching [Pl]_eq_. In the Methods section, we outline quantitative analysis of this survival scenario, which confirms that the qualitative outline presented above indeed explains the persistence of plasmids in some cell lineages despite the ongoing CRISPR interference. Our analysis is based on numerical solution of a master equation that describes time evolution of the probability P_n_(t) for a cell to have n plasmids at time t. The master equation accounts for plasmid replication and interference processes, which are assumed to follow the Logistic dynamics and Michaelis-Menten kinetics, and the binomial partition of plasmids between daughter cells upon mother cell division. The results of the master equation solution for two initial conditions, a cell with a plasmid initially present in a single copy or a cell with a stationary PCN [Pl]_st_ = 100 are presented in Fig. 6e,f. The competition between interference and plasmid replication produces two cell subpopulations, one having a substantial number of plasmids distributed around [Pl]_eq_, and another completely devoid of plasmids. The probability for a cell to retain plasmids quickly drops in the first few generations and levels after 5-10 generations. It follows from our model that the fraction of cells that lose (and, reciprocally, retain) plasmids after this initial transitory period depends on the initial plasmid number in a cell with active CRISPR system. However, the distribution of PCN in cells that retain plasmids converges after ~10 generations to a universal form, which does not depend on the initial number of plasmids and is determined solely by the kinetics of interference and replication. The universal distribution is shown by pale blue lines in Fig. 6e for a single initial plasmid per cell and in Fig. 6f for [Pl]_st_ = 100 plasmids per cell (the fraction of cells with plasmids is much larger for the case with multiple initial plasmids (Fig. 6f) than when there is a single plasmid at the beginning of the process (Fig. 6e), hence the pale blue line in Fig. 6f is higher than that in Fig. 6e).

Since the average PCN converges to a rather large number [Pl]_eq_ (which is independent of the initial conditions), the subsequent probability to lose all plasmids becomes quite low, and such cells with plasmids form colonies that survive indefinitely on antibiotic medium. Thus, our model recapitulates three key experimental observations:

- Under pressure from CRISPR-Cas initially clonal cellular populations become bimodal, consisting of the subpopulations with and without plasmids.
- The fraction of cells that retain plasmids is affected by the distribution of plasmids that exists before CRISPR interference commences.
- The distribution of plasmids under pressure from CRISPR-Cas depends solely on the nature of CRISPR-Cas and plasmids, rather than on the initial plasmid distribution.

Obviously, the model fidelity can be improved by utilizing experimentally-derived dependencies of rates vs. PCN, or fitting the Michaelis-Menten and Logistic constants to the experimental data. However, we believe that the shape or the rate curves illustrated in Fig. 6d is universal. Since the stochastic survival of plasmids goes “against the odds” dictated by an excess of the CRISPR interference rate over the plasmid replication rate, the fraction of cells that retain plasmids falls dramatically with an increase of the rate-reversal threshold [Pl]_bif_ (shown in Fig. 6d-f by a left vertical line). So a fairly delicate balance between the Michaelis-Menten and plasmid replication rate constants, which determine [Pl]_bif_, is required to observe the reported plasmid survival in a small fraction of cells. Within our model, at a single-cell level, the outcome of the plasmid-CRISPR conflict is purely random and all that can be predicted for a given cell is its probability to lose all or retain a certain number of plasmids. A possible determinant of fate of plasmids in a cell could be the level of Cas proteins and plasmid replication machinery enzymes, which themselves are fluctuating quantities. Evidently, cells with above average concentration of plasmid replication machinery enzymes and below average concentration of Cas complexes will have a higher probability to retain plasmids, and vice versa. A “double-stochastic” model that takes into account not only the randomness of replication and interference events, but also fluctuations in levels effector complexes and plasmid replication machinery components could provide even more realistic predictions.

A common way to escape CRISPR interference by phages is the acquisition of mutations in targeted protospacer or its PAM, which decreases and/or abolishes effector complex affinity^6,26^. Indeed, phage plaques formed on lawns of cells identical to the ones used here with CRISPR targeting various phages are formed by such escaper phages^27^. Yet, no escape plasmids are found in colonies formed upon transformation under CRISPR ON conditions^14^, even though escape plasmids created in the laboratory are efficiently transformed and are not subject to CRISPR interference^4,6^. We hypothesize that the difference between phage and plasmid reactions to an ongoing CRISPR interference is due to the fact that the former but not the latter are cell-autonomous in a sense that they can be released from the infected cell and then reinfect surrounding cells. Thus, during formation of a plaque (a negative colony) multiple reinfections take place which allows rare phages that acquire escape mutations to take over the population, such that the final plaque contains almost exclusively mutant phages^6^. In contrast, during formation of a bacterial colony by a founder cell transformed with a plasmid, plasmids are only passed vertically from parent to daughter cells. While an infected cell usually receives just one copy of the phage genome, a daughter cell inherits on average half of plasmids from its mother. Hence, if the probability for a phage to survive and replicate in a cell (which is quite low in the case of a single initial plasmid considered here), multiplied by the size of phage burst is much smaller than one, the infection would not propagate and we would simply register an apparent defeat of a phage by CRISPR-Cas.

A limited rate of CRISPR interference resulting in an incomplete extermination of phages and plasmids can be a consequence of simple evolutionary principles. Increasing the rate of interference costs the cell not only extra energy to produce additional copies of Cas proteins but can also lead to off-target DNA cleavage, causing autoimmunity. Thus, the evolutionary optimization of CRISPR-Cas interference rate would probably not go beyond some intermediate protection level, which eliminates foreign mobile genetic elements in most but not in every cell in the population. The stochasticity of CRISPR interference could allow conditionally favourable plasmids to take a hold in a subpopulation of cells and then proliferate when the environment selects^28^. Such effects can be especially prominent in structured environments offering specific niches to subpopulations differing in their genetic or physiological states^29,30^. Bacterial colonies and biofilms are not homogeneous and contain micro-environments with complex spatial structures^31,32^ that may support, among other things, cooperative antibiotic^22,23,33^ or phage resistance^34,35,36^. Some of such complex spatial structures may be similar to those microscopically observed in CRISPR ON colonies in our experiments and depend on a dynamic interplay between mobile genetic elements replication and host defence directed against them.

## Methods

### Strains and plasmids

*E. coli* strain KD263 (K-12 F+, *lac*UV5-*cas3* araBp8-*cse1*, CRISPR I: repeat-spacer g8-repeat, CRISPR II deleted) has been described^17^. The pG8 plasmid carrying a 209-bp phage M13 fragment with the g8 protospacer has been described^11^. The pRSFG8 plasmid carrying the 209-bp phage M13 fragment with the g8 protospacer has been constructed previously^18^. The pG8-GFP plasmid was derived from pG8 by cloning the *TagGFP2* gene (Evrogen) following the Gibson assembly protocol (NEB). Primers used for DNA amplification are listed in Extended Data Table 2. *E. coli* cells were grown at at 37 °C in LB medium (per 1L: 5g NaCl, 10 g tryptone, and 5 g yeast extract) or on LB-agar plates containing 1.5% agar.

### CRISPR interference assay

*E. coli* strain KD263 overnight culture was diluted 100 times into 5 mL of LB. The cells were grown in the presence (CRISPR ON) or in the absence (CRISPR OFF) of 1 mM arabinose and 1 mM IPTG for *cas* genes expression until cultures OD_600_ reached 0.6. The electrocompetent cells were prepared following a standard protocol^37^ and transformed with 5 ng of the protospacer plasmid (pG8 plasmid or pRSFG8). Next, the transformed cells were grown in 1 ml of LB supplemented with 1 mM arabinose and 1 mM IPTG for CRISPR ON cultures and 1 ml of LB for CRISPR OFF cultures for 1 h. The 50 μl aliquots of serial dilutions of the transformation mixtures were plated onto LB agar plates containing 100 μg/ml ampicillin (for pG8 plasmid transformation) or 50 μg/ml kanamycin (for pRSFG8 plasmid transformation) (CRISPR ON) or without (CRISPR OFF) inducers. The plates were incubated at 37 °C overnight. The efficiency of transformation (EOT) was determined as a number of colony forming units (CFU) per 1 μg of plasmid DNA (Fig. 1a,b). Each transformation was performed in triplicate.

To test the condition of the protospacer plasmids in CRISPR ON transformants, plasmid DNA from ten randomly chosen and pooled CRISPR ON colonies was isolated using GeneJET Plasmid Miniprep Kit (Thermo scientific) and retransformed into fresh prepared CRISPR ON and CRISPR OFF cells (Fig. 1c). To test the functionality of CRISPR-Cas system in CRISPR ON transformants, retransformation of cells derived from ten randomly chosen individual CRISPR ON colonies was carried out with the second plasmid bearing the compatible origin and the g8 protospacer: the cells initially transformed with pG8 plasmid received pRSFG8 plasmid and vice versa (Fig. 1d). The efficiency of transformations was determined as described above.

### Quantitative PCR assay of plasmids

qPCR was performed using DTlite4 Real-Time PCR System (DNA-Technology). Reactions were carried out in triplicate (technical repeats) in a 20 μl reaction volume supplemented with 0.8 units of HS Taq DNA polymerase, 2 μl 10x Taq Turbo buffer (Evrogen), 0.25 mM dNTPs, 0.2 μl Tween 20, 0.1 μl of SYTO™ 13 intercalating dye (LifeTechnology), 1 μl of sample and appropriate primers at 5 pM. The primers for qPCR are listed in Extended Data Table 2. Three randomly chosen colonies were suspended in 20 μl distilled water. The results of qPCR with plasmid-specific primers were normalized to genomic DNA with regard to efficiency of the primers (Supplementary Data 1).

The efficiency of the primers was calibrated following a standard curve^38^. To calculate the standard curve for the primers, three random individual colonies of the transformants were chosen and suspended in 20 μl distilled water. Next, three 10-fold serial dilutions of the suspended samples were assayed with qPCR using three technical replicates for each sample. We used only the results of qPCR with the deviation less than 0.1 ΔCt (cycle threshold) among technical repeats of one dilution. The efficiency of primers was calculated as an average slope of the plot of logarithm of the concentration of the dilutions vs. ΔCt (Supplementary Data 1). Three repeats for each group of the primers were carried out. The following efficiencies of the primers were obtained: 2.0 for Bla_dir and Bla_rev (amplification efficiency 100%), 1.94 for GyrA_dir and GyrA_rev (amplification efficiency 94%) and 2.08 for pRSF_ori_dir and pRSF_ori_rev (amplification efficiency 108%). Mean PCN was estimated as a ratio of genomic to plasmid ΔCt values considering the efficiency of the primers (Supplementary Data 1).

### Replating of transformants

Four randomly chosen individual colonies of the CRISPR ON and CRISPR OFF transformants were replated on three types of selective media: LB supplemented with appropriate antibiotic (Ab) and inducers (Ind) to maintain the CRISPR-Cas activity, LB supplemented with appropriate Ab only (to determine the number of plasmid-bearing cells) and LB (to determine the total number of cells) (Fig. 2a). Each colony was suspended in 500 μl of LB and eight 4-fold serial dilutions of the suspended cells were prepared. Next, 5 μl of each dilutions were plated on the selective media. The CFUs were counted on each plate. The colonies from plates with Ab/Ind were used for the second replating. Each replating was repeated at least three times.

### Flow cytometry

Several colonies of *E. coli* CRISPR ON and OFF cells bearing the pG8-GFP plasmid were suspended in PBS (phosphate-buffered saline) and passed through 100-μm filters. Samples were investigated using FACSAria III (BD Biosciences); the flow cytometry protocol was customized for bacterial cells. Forward versus side scatter (FSC vs SSC) plots were used to gate the area of single cells; 2 × 10^5^ events per sample in the gate was collected. TagGFP2 fluorescence was excited with 488-nm laser and detected with 530/30 filter. Three biological replicates for each sample were done. The data were analysed by FCSalyzer and Flowing Software.

### Microscopy assay

Fluorescence imaging microscopy of CRISPR OFF and ON colonies was performed using Leica AF6000 LX system based on a DMI 6000 B inverted microscope equipped with HCX PL APO lbd. BL 63x 1.4NA oil objective and Photometrics CoolSNAP HQ CCD camera. GFP filter cube (excitation BP470/40 and emission BP525/50) was used to visualize TagGFP2. LB-agar fragments containing colonies were cut, placed on glass bottom dishes so that the colonies were adjacent to the glass bottom. The colonies were observed in fluorescence and transmitted light channels. Colonies of *E. coli* KD263 cells transformed with pG8 were used as a negative control (no fluorescent protein). CRISPR OFF and ON colonies were also visualized with laser scanning confocal microscope DMIRE2 TCS SP2 (Leica) with HCX PL APO lbd.BL 63x 1.4 NA oil objective. The green fluorescent signal was acquired at 488-nm excitation and detected at 500-to 530-nm wavelength range.

### Single cell microscopy in microfluidics device

#### Design of microfluidics device

The device was designed using AutoCAD^®^ (AUTODESK^®^) and the Metafluidics database and fabricated from PDMS following standard soft lithography technique^39^. It includes four major trenches of 100 μm width and 40 μm depth each, along which the growth medium is passed, and 1000 growth chambers with the depth of about 1 μm and the length of 20 μm on the front side that adjacents to the major trench and 80 μm on the lateral side. The inlet of the device contained a 25 μm filter to prevent clogging. To make the device a double layer mold was fabricated using SU - 8 2025 photoresist (Kayaku Advanced Materials, Inc) spin coated onto a silicon wafer and exposed by contact photolithography with two chromium masks. For the first layer SU - 8 2025 was diluted by SU - 8 T thinner to achieve the thickness of the layer about 1 μm. After that the PDMS prepolymer and the curing agent (Sylgrad 184, Dow Corning) were mixed in a ratio of 10:1 w/w, degassed, poured into the mold, and cured at 65 °C for 4 h in an oven. Then the PDMS replica was detached from the mold, inlet and outlet holes were made by a 1 mm biopsy puncher. Finally the replica was bonded with a cover glass slide after oxygen plasma treatment.

#### Single cell microscopy

Several colonies of *E. coli* CRISPR OFF cells bearing pG8-GFP plasmid were resuspended in LB medium with 1 mM arabinose and 1 mM IPTG inducers for *cas* genes expression and loaded to the microfluidics device. Single fluorescent cells caught in the growth chambers were tracked using a Nikon Eclipse Ti-E inverted epifluorescence microscope. Cells were cultivated in the growth chambers overnight at 37 °C on LB medium supplemented with *cas* genes expression inducers. The images were captured every 15 minutes for generating a time-lapse movie in transmitted light for all cell observation and in the green channel for fluorescence detection. The tagGFP2 fluorescence signal was detected using the Semrock filter set YFP-2427B. All images were obtained using Zyla 4.2 sCMOS camera (Andor). Fluorescence intensity from single cells was analysed using ImageJ software.

### Dynamics of replication and degradation of plasmids

The dynamics of plasmid replication can be quite complex, yet it has two universal limits: For a few plasmid, the replication rate is proportional to the number of plasmids (i.e. the replication rate per plasmid is constant), while for the target (target) concentration of plasmids [Pl]_*st*_, the replication rate is zero. As it is often done^39^, we approximate such dynamics by the Logistic model,

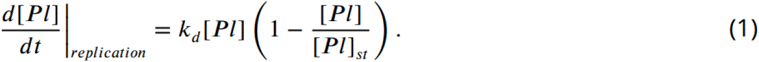

The coefficient *k_d_* is the per capita rate of plasmid replication in the low concentration limit. The symbol [*x*] indicates the concentration of a substance *x*. Assuming that the volume of a cell stays approximately constant, we define a concentration as the number of molecules per cell, and in the following we use the terms “concentration” and “copy number” interchangeably.

As a catalytic process, the interaction of CRISPR-Cas complexes *Cr* with plasmids *Pl*,

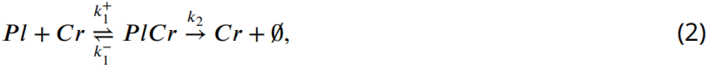

is assumed to be well-described by the Michaelis-Menten kinetics,

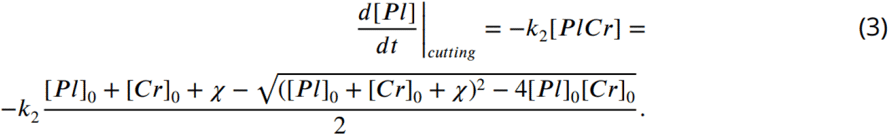

Here, as in the standard Michaelis-Menten derivation, the stationarity of concentration of the CRISPR-Cas-plasmid complex is assumed, the generalized dissociation constant *χ* is defined as

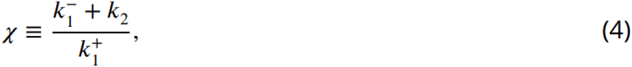

and no assumption is made about overabundance of the catalyst (CRISPR-Cas) or the substrate (plasmid). The total (bound in the *Pl-Cr* complex and free) concentrations of plasmids and CRISPR-Cas complexes are denoted as [Pl]_0_ and [Cr]_0_.

Assuming that replication only increases the plasmid concentration so that [Pl] in (1) never exceeds [Pl]_st_, we define a one-step birth-death process^41^ for the population of plasmids. The probabilities of increasing or decreasing the population of plasmids by one *β*_[Pl]_ and *δ*_[Pl]_ are given by *d*[Pl]/*dt*|_replication_ (1) and *d*[Pl]/*dt*|_cutting_ (3). The master equation that describes the temporal evolution of probability *P*_[Pl]_ (*t*) to find a cell having [Pl] plasmids at time *t*^41^ reads

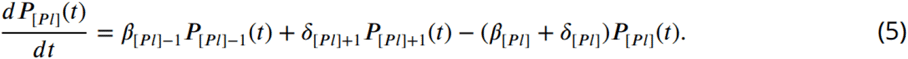

### Redistribution of plasmids during cell division

In addition to cutting and replication of plasmids, the per cell PCN is also affected by cell division, which on average happens every *τ* ≈ 20 min. A conservative estimate would be that the redistribution of plasmids between daughter cells is completely random (in reality it is biased towards equal or half and half distribution). Assuming also that the act of cell division happens fast (instantaneous) compared to the replication and cutting of plasmids, the outcome of the redistribution process can be described by the binomial distribution with the probability for each plasmid to go into any of two daughter cells equal to 1/2. If a cell before the division had *j* plasmids, then the probability *B*_ij_ to find 0 ≤ *i* ≤ *j* plasmids in one of the daughter cells is

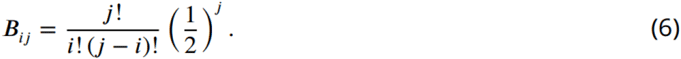

### Simulation procedure

As presented above, the temporal dynamics of plasmid copy number in a cell is approximated by a sequence of periods of continuous evolution, described by the master equation (5), each followed by the instantaneous redistribution between daughter cells, described by the binomial distribution (6). To estimate the distribution of plasmids in cells in CRISPR ON colonies after several hours of growth, we implement the following numerical procedure:

- For a given set of plasmid replication and CRISPR interference parameters *k_d_*, [Pl]_st_, *k_2_, χ*, and [Cr]_0_, we tabulate the replication and cutting rates *β*_[Pl]_ and *δ*_[Pl]_ for all possible plasmid copy numbers, 1 ≤ [Pl] ≤ [Pl]_st_.
- We numerically integrate the master equation (5) till the cell cycle time *τ*, starting from every possible initial number of plasmids *j*, 0 ≤ *j* ≤ [Pl]_st_. Naturally, the solution with zero initial plasmids will always be zero plasmids with probability one.
- The probabilities *C*_ij_ for a cell to end up with *i* plasmids at time *τ* after starting with *j* plasmids at *t* = 0,

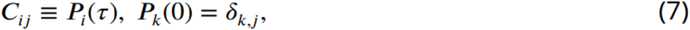

are collected into the matrix *Ĉ*. Another matrix 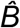 is composed of binomial probabilities *B*_ij_ (6).
- The probability to find *k* plasmids after time *t* is given by the *k* + 1th element of the [Pl]_st_ + 1-dimensional vector 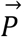,

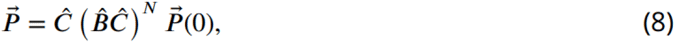

where *N* (equal to the integer part of *t*/*τ*) is the number of cell cycles and the initial condition 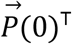 indicates how many plasmids were in each cell when the CRISPR-Cas system was activated. Here we assumed that the number of plasmids in a cell is assessed at the final stage of cell cycle just before cell division, thus an extra multiplication by *Ĉ*. (Alternatively, when the number of generations is not very large, this probability can be computed more efficiently by direct solution of the master equation (5) for periods of time between cell divisions, alternated with binomial redistribution of plasmids between daughter cells according to (6). In such a case we do not need to compute the matrix *Ĉ*).

The evolution of the probability density *P_k_*(*t*) for the replication and interference rates (1) and (3) plotted in Fig. 6d is shown in Fig. 6e for cells initially having 1 plasmid, (*P_k_*(0) = *δ*_k,1_ being the typical initial condition in a CRISPR-ON experiment) and in Fig. 6f for cells initially having the target number of plasmids, (*P_k_*(0) = *δ*_k,[Pl]st_ being the initial condition for replating the CRISPR OFF cells on plates with inductor). The plots in Fig. 6 were computed using the following parameters *k*_d_ = 0.3, [Pl]_st_ = 100, *k*_2_ = 0.5, *χ* = 1, and [Cr]_0_ = 10.

As many birth-death processes, this stochastic process of plasmid replication, cutting, and redistribution has the unique convergent steady state 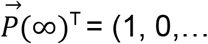, corresponding to the extinction of all plasmids. However, after a few cell cycles, while the component *P*_0_(t) that corresponds to the fraction of cells with no plasmids steadily grows, other components *P*_k_(t), *k* = 1,…, *k_st_* that correspond to the probability to have a non-zero number of plasmids approach a steady state scaling form,

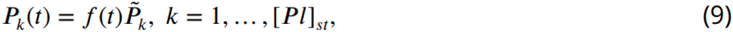

shown in Extended Data Fig. 6. The slowly-decaying function *f*(t) represents a universal convergence to the absorbing state 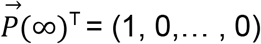.

## Supporting information

https://drive.google.com/file/d/1st1lSLlluNDIstWJpfkxvMOhVLcVzrZ6/view?usp=sharing

## Acknowledgments

We thank A. Strotskaya for providing plasmids. This study was supported by NIH grant GM10407 to K.S. Y.I. is supported by FONDECYT project 1200708. Experiments were partially carried out using the equipment provided by the Institute of Bioorganic Chemistry Core Facility (CKP IBCH; supported by Russian Ministry of Education and Science Grant RFMEFI62117X0018 to K.L.)

## Extended Data

**Extended Data Figure 1.**
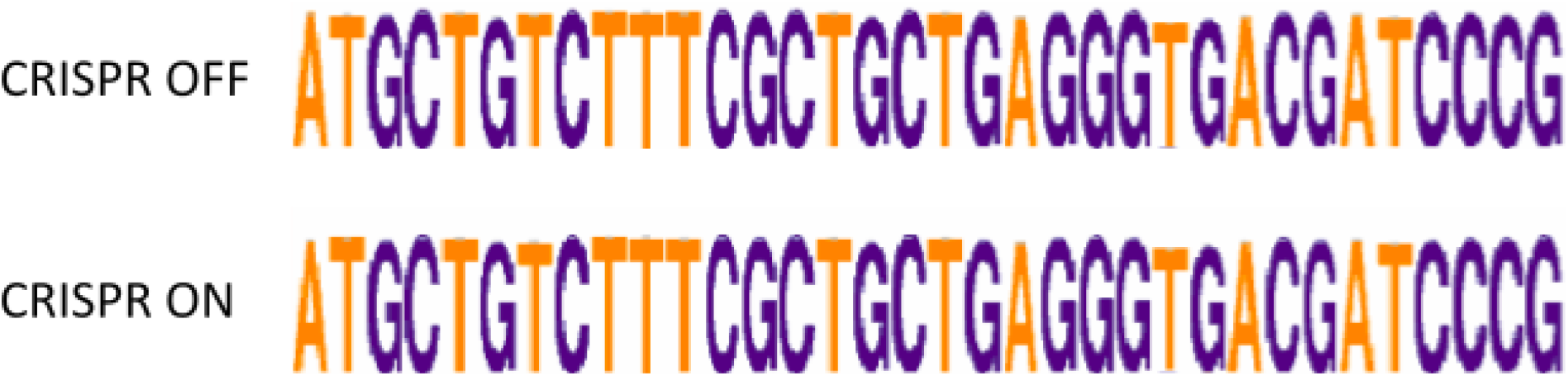
Sequences of PAM and g8 protospacer derived from a pool of pG8 plasmids. Sequence logos show no escape mutations in PAM and protospacer of the pG8 plasmid purified from CRISPR ON cells similar to the plasmids purified from CRISPR OFF cells.

**Extended Data Figure 2.**
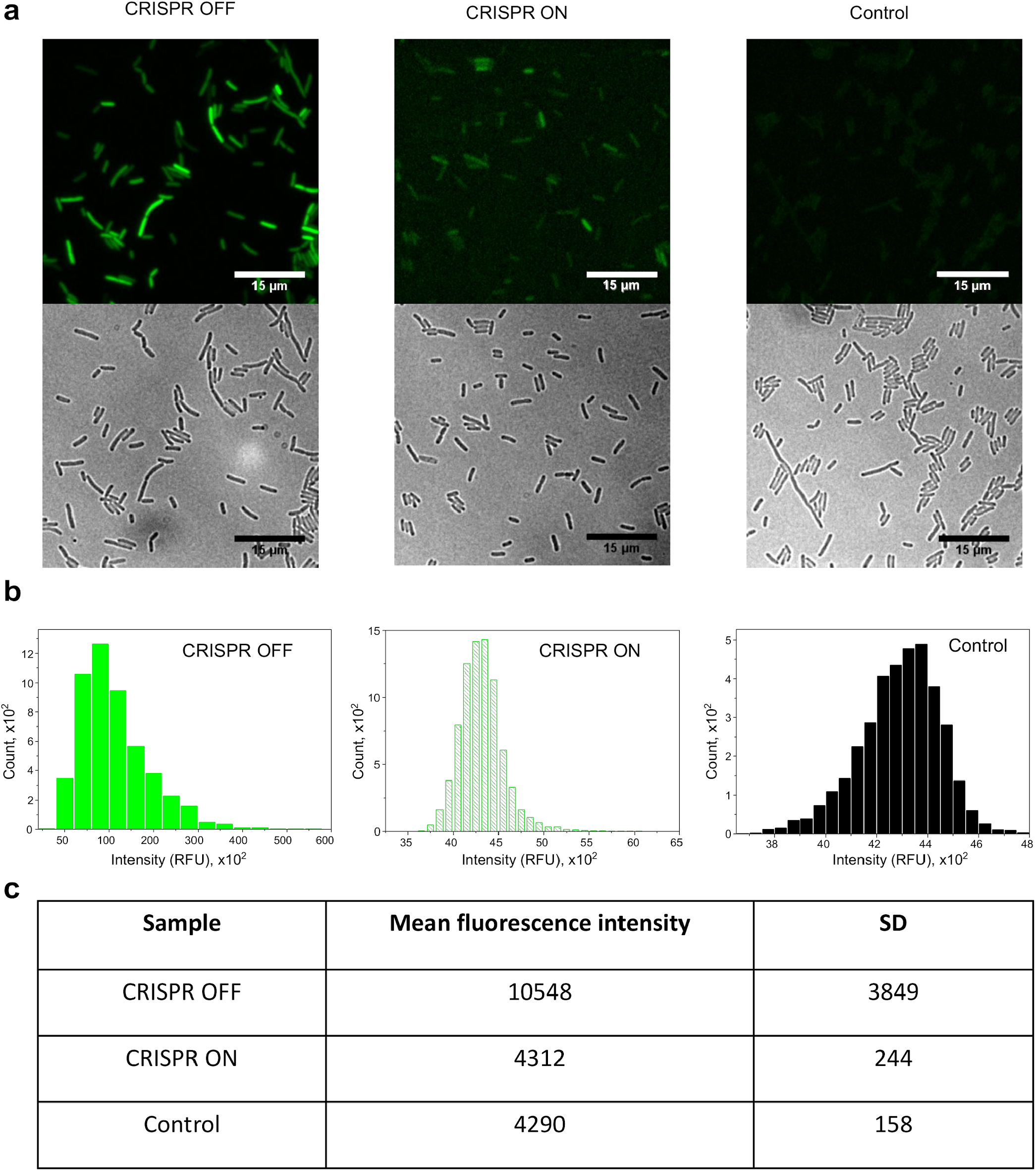
Fluorescence intensity of CRISPR ON and CRISPR OFF transformant cells. **a,** Images of cells derived from CRISPR ON and CRISPR OFF colonies. KD263 cells were used as a negative control. **b,** Histograms show the distribution of fluorescence of cells in relative fluorescence units (RFU). **c,** Statistics of fluorescence intensity.

**Extended Data Figure 3.**
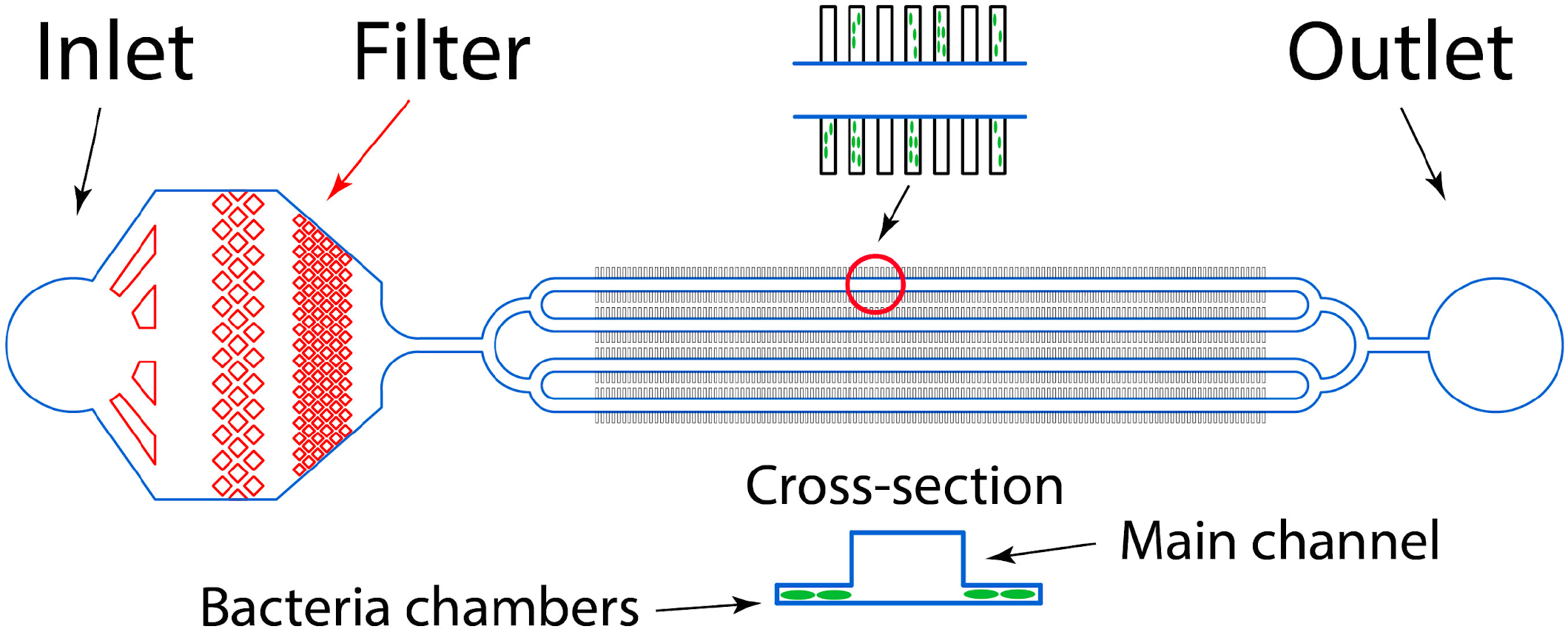
A scheme of the microfluidic device for live cell microscopy. The device contains 1000 growth chambers 80×20×1 μm located on both sides of the four major channels of 100 μm width and 40 μm depth each that are used for introducing bacteria cells into the growth chambers and for the fresh LB medium circulation. The inlet of the device contains a 25 μm filter to prevent clogging.

**Extended Data Figure 4.**
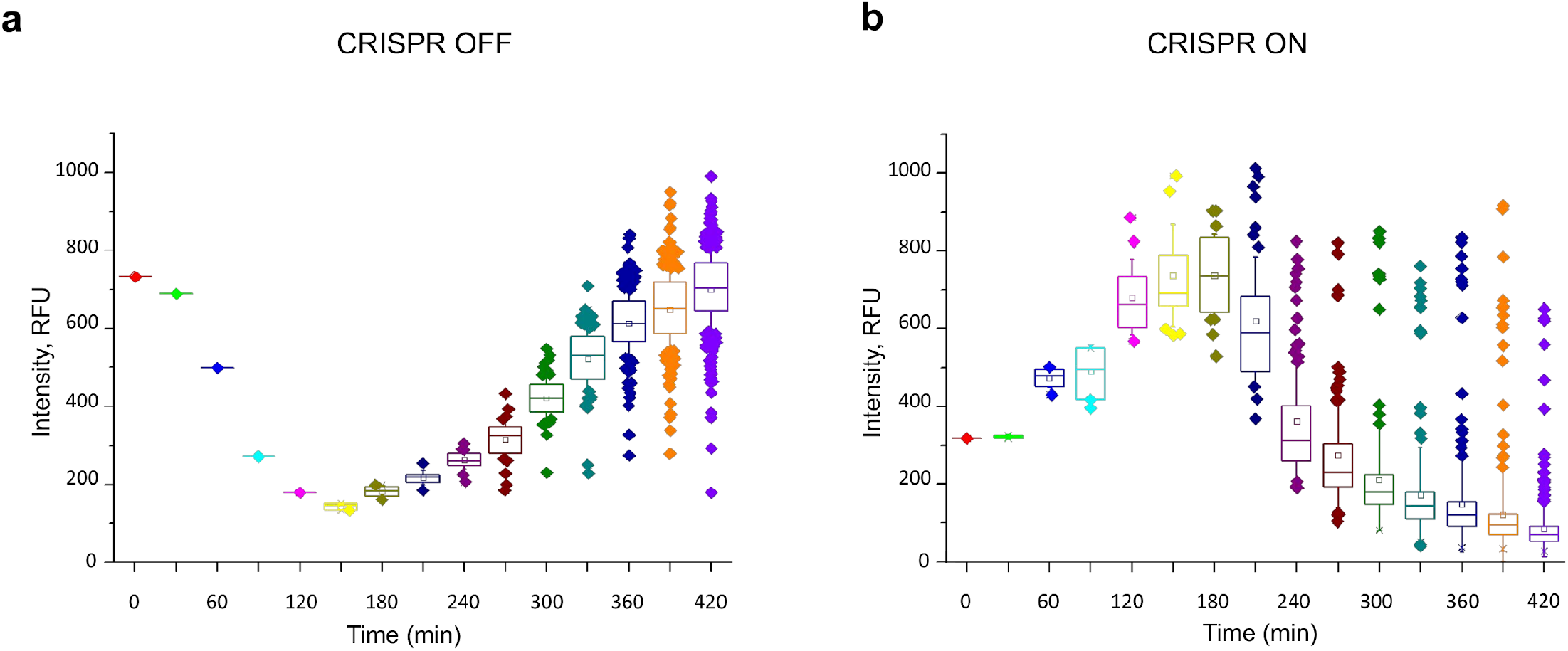
Fluorescence intensity change of CRISPR ON and CRISPR OFF cells in microfluidic channels over time. **a,** CRISPR OFF fluorescent cells show normal fluorescence intensity in relative fluorescence units (RFU) as a positive control. **b,** The fluorescence intensity of cells in CRISPR ON experiment decreases over time after activation of the CRISPR-Cas system at 2-3 hour points, but a small amount of cells (≈6-7%) remain fluorescent. At the same time, the intensity of fluorescent CRISPR ON cells becomes less than the mean level (≈700 RFU) inherent to CRISPR OFF cells.

**Extended Data Figure 5.**
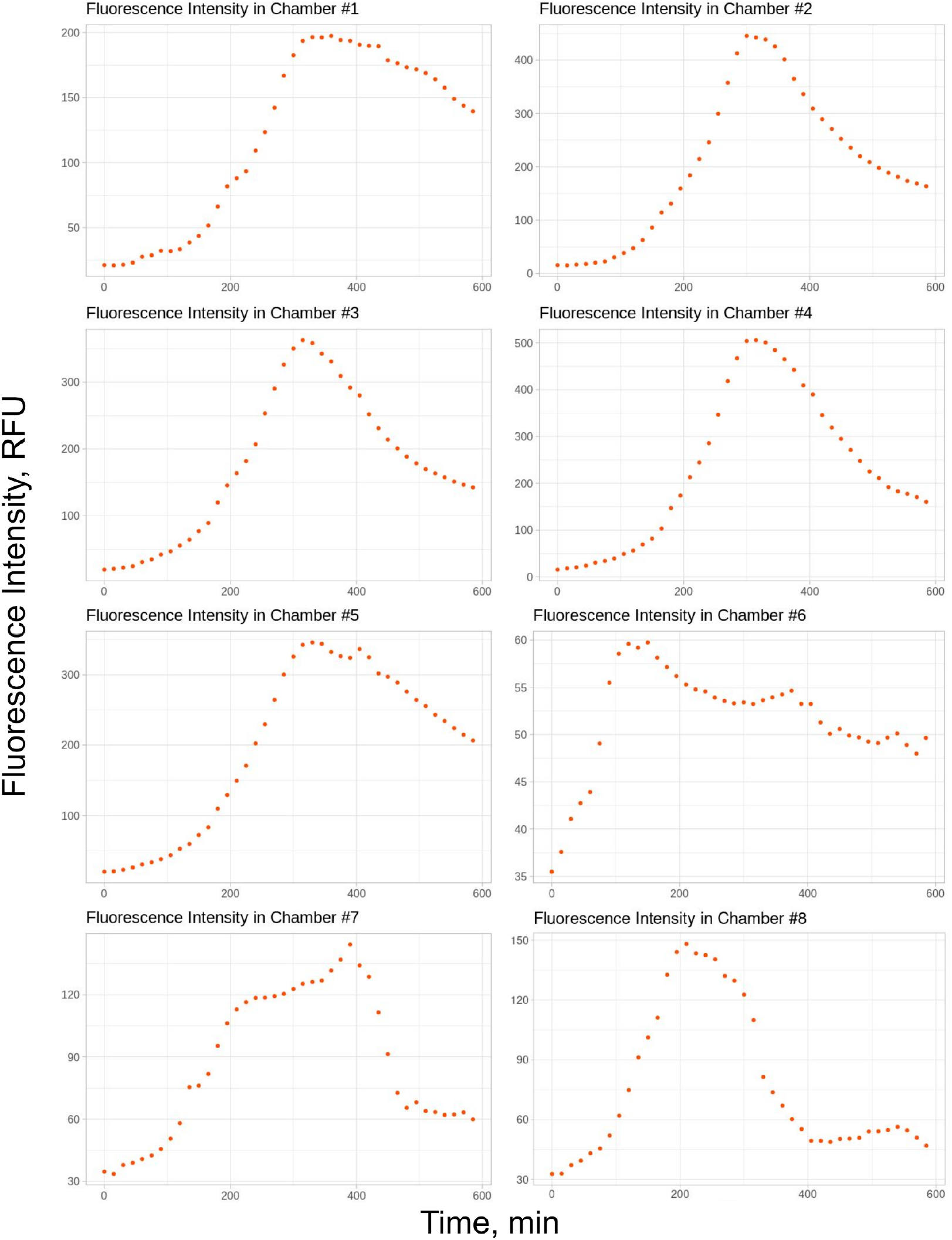
Mean fluorescent intensity per microfluidics channel area over time. Cultivation of KD263 cells bearing plasmid pG8-GFP was observed in LB media supplemented with *cas* genes expression inducers. Mean fluorescence intensity in relative fluorescence units (RFU) from all KD263 cells for each growth chamber was normalized on background level and calculated. All plots demonstrate an increase of mean fluorescence intensity at the initial stages of observation, but starting from 300 min after induction the fluorescence intensity decreases.

**Extended Data Figure 6.**
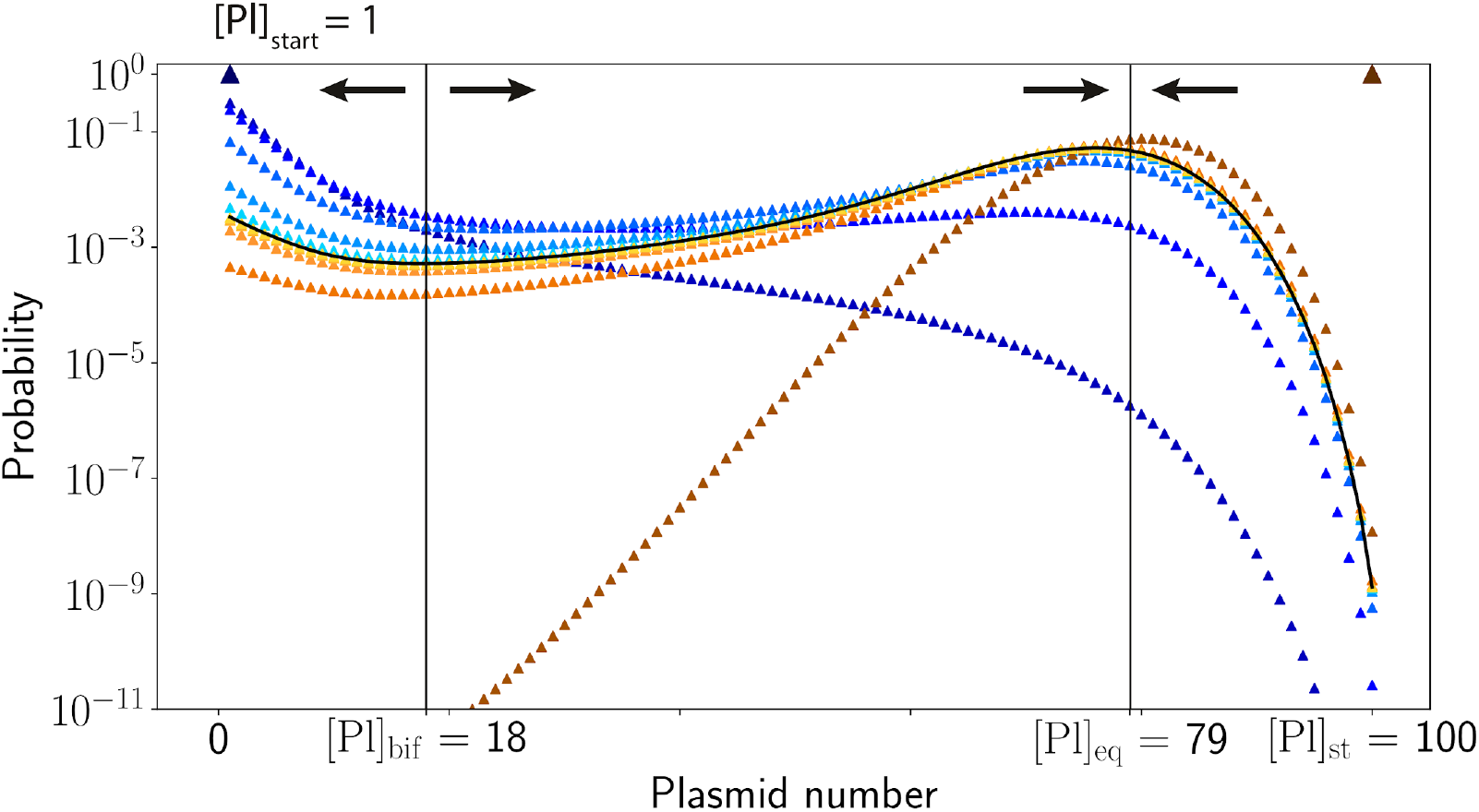
Convergence of PCN probability distribution to the universal scaling form. The blue triangles show the evolution of *P_k_*(*t*) when cells had initially a single plasmid, while the orange/brown squares show the evolution of *P_k_*(*t*) when cells had initially [*Pl*]_st_ plasmids. Shades of blue and red correspond to the different generations of cells from generation 0 (dark triangle and square) to generation 5 (the lightest blue and orange). Both families of curves converge to the universal asymptotic curve shown by a black line.

**Extended Data Table 1.**
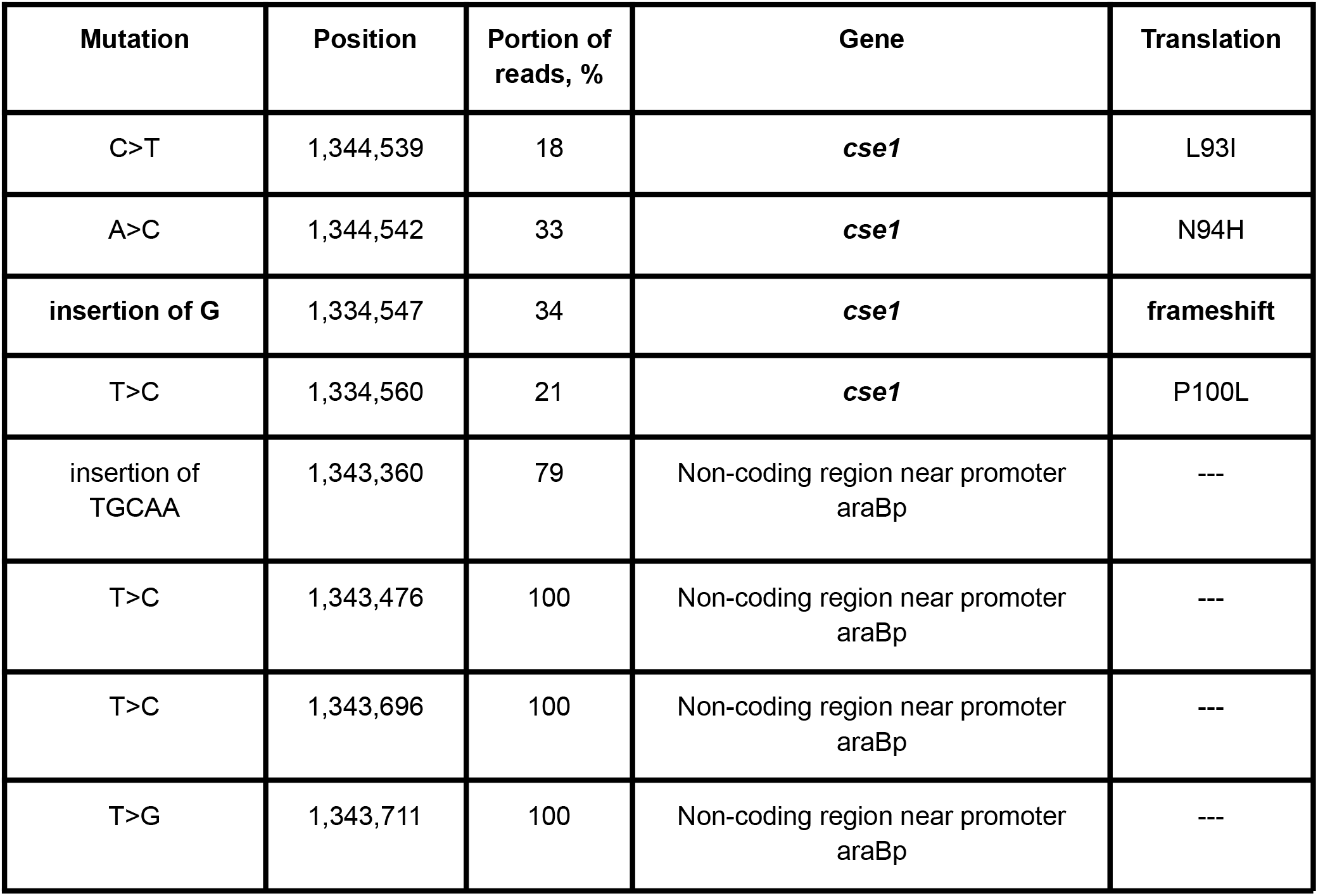
The list of mutations disrupting CRISPR-Cas activity in cells from bright CRISPR ON colonies.

| **Mutation** | **Position** | **Portion of reads, %** | **Gene** | **Translation** |
| --- | --- | --- | --- | --- |
| C>T | 1,344,539 | 18 | ***cse1*** | L93I |
| A>C | 1,344,542 | 33 | ***cse1*** | N94H |
| **insertion of G** | 1,334,547 | 34 | ***cse1*** | **frameshift** |
| T>C | 1,334,560 | 21 | ***cse1*** | P100L |
| insertion of TGCAA | 1,343,360 | 79 | Non-coding region near promoter<br>araBp | --- |
| T>C | 1,343,476 | 100 | Non-coding region near promoter<br>araBp | --- |
| T>C | 1,343,696 | 100 | Non-coding region near promoter<br>araBp | --- |
| T>G | 1,343,711 | 100 | Non-coding region near promoter<br>araBp | --- |

**Extended Data Table 2.**
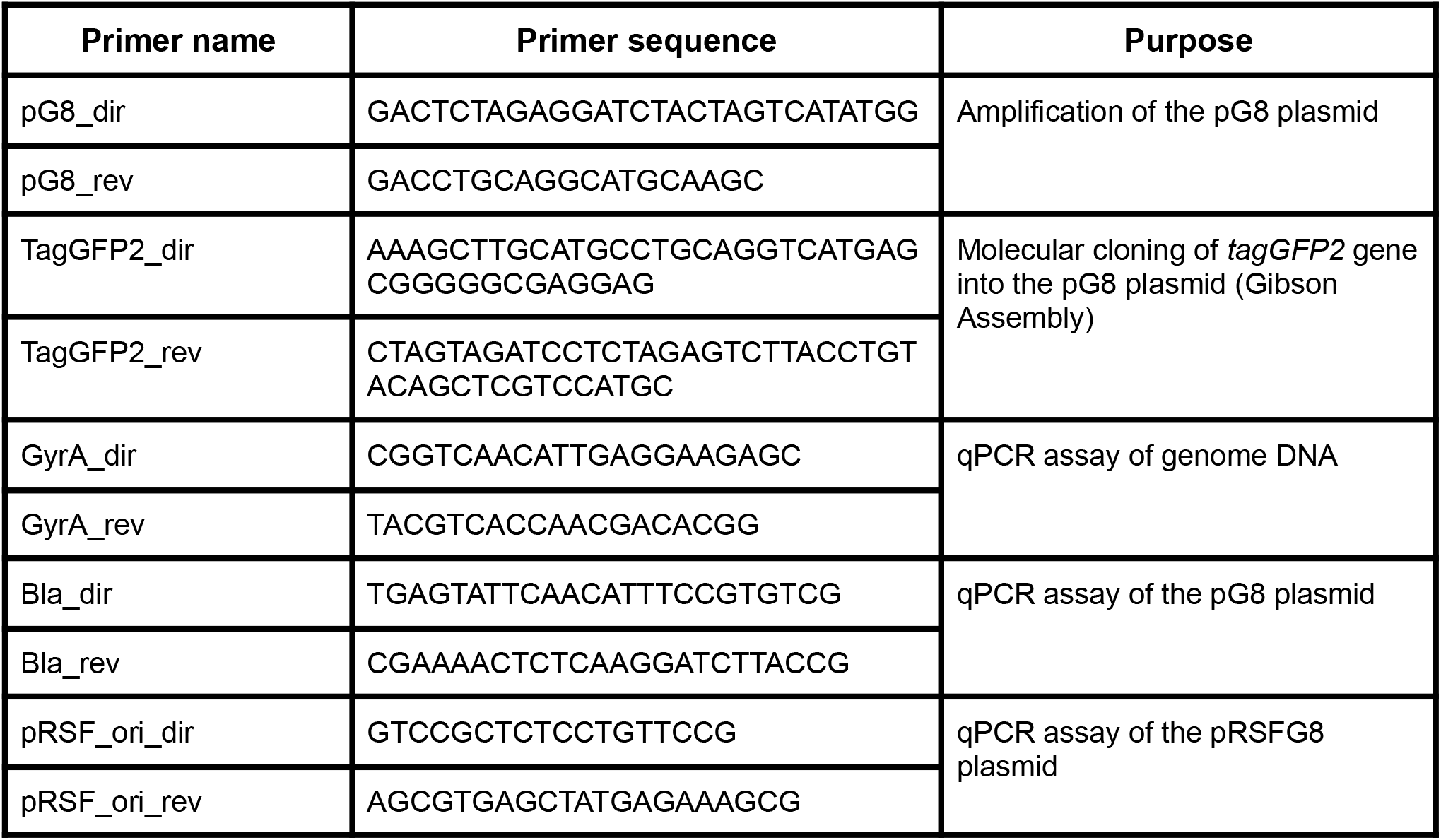
Nucleotide sequences of primers used for cloning.

